# Drug-resistant epimutants exhibit organ-specific stability and induction during murine infections caused by the human fungal pathogen *Mucor circinelloides*

**DOI:** 10.1101/636050

**Authors:** Zanetta Chang, Joseph Heitman

## Abstract

The environmentally ubiquitous fungus *Mucor circinelloides* is a primary cause of the emerging disease mucormycosis. *Mucor* infection is notable for causing high morbidity and mortality, especially in immunosuppressed patients, while being inherently resistant to the majority of clinically available antifungal drugs. A new, RNAi-dependent, and reversible epigenetic mechanism of antifungal resistance – epimutation - was recently discovered in *M. circinelloides*. However, the effects of epimutation in a host-pathogen setting were unknown. We employed a systemic, intravenous murine model of *Mucor* infection to elucidate the potential impact of epimutation *in vivo*. Infection with an epimutant strain resistant to the antifungal agents FK506 and rapamycin revealed that the epimutant-induced drug resistance was stable *in vivo* in a variety of different organs and tissues. Reversion of the epimutant-induced drug resistance was observed to be more rapid in isolates from the brain, as compared to those recovered from the liver, spleen, kidney, or lungs. Importantly, infection with a wild-type strain of *Mucor* led to increased rates of epimutation after strains were recovered from organs and exposed to FK506 stress *in vitro.* Once again, this effect was more pronounced in strains recovered from the brain than from other organs. In summary, we report the rapid induction and reversion of RNAi-dependent drug resistance after *in vivo* passage through a murine model, with pronounced impact in strains recovered from brain. Defining the role played by epimutation in drug resistance and infection advances our understanding of *Mucor* and other fungal pathogens, and may have implications for antifungal therapy.

**IMPORTANCE:** The emerging fungal pathogen *Mucor circinelloides* causes a severe infection, mucormycosis, which leads to considerable morbidity and mortality. Treatment of *Mucor* infection is challenging because *Mucor* is inherently resistant to nearly all clinical antifungal agents. An RNAi-dependent and reversible mechanism of antifungal resistance, epimutation, was recently described in *Mucor*. Epimutation has not been studied *in vivo* and it was unclear whether it would contribute to antifungal resistance observed clinically. We demonstrate that epimutation can be both induced and reverted after *in vivo* passage through a mouse model; rates of both induction and reversion are higher after brain infection than after infection of other organs (liver, spleen, kidneys, or lungs). Elucidating the roles played by epimutation in drug resistance and infection will improve our understanding of *Mucor* and other fungal pathogens, and may have implications for antifungal treatment.

## INTRODUCTION

The fungal infection mucormycosis is caused by a group of related fungi, the most common of which are the genera *Rhizopus, Mucor,* and *Lichtheimia* (1). Mucormycosis encompasses a broad range of infections, from cutaneous infections to invasive systemic disease, which can lead to rates of mortality greater than 90% (2, 3). The most common manifestation of disease is rhino-orbito-cerebral mucormycosis, which is often seen in patients with predisposing factors such as immunosuppression or diabetes (4). Rates of mucormycosis are increasing worldwide, due in part to an increase in the prevalence of predisposing factors (4, 5). As the causative fungi are ubiquitous in the soil, outbreaks are often caused by exposure to environmental sources, ranging widely from environmental disasters such as tornadoes to contaminated hospital linen (6-8). Mucormycosis is highly drug resistant and only three antifungal drugs are approved for treatment: amphotericin B, isavuconazole, and posaconazole (9, 10). Therefore, a better understanding of drug resistance in mucormycosis is necessary for improving clinical outcomes of this emerging pathogen.

Among the Mucorales, the genus *Mucor* is unique in that it also serves as a genetic model. In addition to its pathogenic nature, it has been utilized to study aspects of fungal biology as diverse as biofuel production, RNA interference (RNAi), and light sensing (11-16). It also serves as a model for a novel mechanism of intrinsic, transient, and RNAi-dependent antifungal resistance known as epimutation. In epimutation, *Mucor* endogenous RNAi pathways are activated and induce silencing of various drug target genes to result in antifungal resistance (17, 18). The mechanism of epimutation requires the core RNAi pathway and competes with an alternative RNA degradation pathway in *Mucor* (19).

Here we studied epimutation-based drug resistance with the antifungal drug FK506 (also known as tacrolimus). FK506 binds the target protein, FKBP12, which is encoded by the *fkbA* gene in *Mucor*. Mutation of FKBP12 or repression via RNAi-based silencing of *fkbA* leads to drug resistance (20). The FKBP12-FK506 complex inhibits the calcium-calmodulin activated protein phosphatase calcineurin, which is required for *Mucor* dimorphism and virulence (21, 22). Wild-type *Mucor* strains grow as yeast when exposed to FK506, whereas resistant strains are blocked in the dimorphic transition and grow exclusively as filamentous hyphae in either the presence or absence of FK506 under aerobic conditions.

We report here the first observations of epimutation affecting antifungal resistance after *in vivo* passage through a systemic murine model of mucormycosis. Infection of mice with an epimutant strain resistant to FK506 revealed that epimutants are largely stable during the course of infection. However, when epimutants were observed to revert during *in vivo* infection, the reversion occurred most prominently in the brain compared to other organs analyzed. Interestingly, infection with a wild-type strain led to an increased induction of epimutant-based resistance when isolates were recovered from *in vivo* passage and exposed to FK506 *in vitro*. Induction of resistance was also observed to be more frequent in fungal isolates recovered from the brain compared to other organs. These observations suggest that rapid reversion and induction of epimutation may enable fungal microbial responses to transitions between the environment and the host. Elucidating the mechanisms by which *Mucor* responds to stressful environmental conditions, including factors as diverse as vertebrate hosts and antifungal drug stress, advances our understanding of mucormycosis, fungal pathogens, and the mechanisms and pathways that lead to the development of antifungal drug resistance and its maintenance and loss.

## RESULTS

### Systemic intravenous infection of mice via retroorbital inoculation

For the murine infections in this work, we developed a model of *Mucor* infection via the retroorbital venous sinus (23). Intravenous infection of mice via tail vein injection of *Mucor* spores is an established model of systemic infection that results in high mortality (24). However, tail vein injections are technically challenging, which can influence reproducibility of experimental results. Therefore, we wished to develop an alternative systemic infection model. The retroorbital route of injection has been established to be safe and effective, and has been shown to be equivalent to tail vein injection for systemic delivery of intravenous drugs; it has also previously been used to deliver larger moieties to the systemic circulation, including monoclonal antibodies and tumor cells (25, 26). In brief, mice are anesthetized with inhaled isoflurane, the area behind either eye is injected with fungal spores in saline solution, and the animal is removed from anesthesia and rapidly recovers over approximately 30 seconds.

To determine whether retroorbital infection would produce the same course of disease as tail vein infection, we compared the two injection methods using groups of 5 BALB/c mice. Spores of the 1006PhL strain of *Mucor circinelloides* f. *circinelloides* were injected into immunocompetent, wild-type mice by either the retroorbital or tail vein methods at a dose of 1.25 × 10^6^ spores/100 μL PBS per mouse. Despite longstanding experimental experience with the tail vein injection method, technical difficulties were experienced when injecting 3 of the 5 mice via tail vein, potentially due to the viscosity of the concentrated spore solution; no issues were noted with any animals in the study group while injecting retroorbitally. All infections were allowed to progress until mice were euthanized at humane endpoints. No significant difference was noted between retroorbital and tail vein groups with regard to time to mortality (Fig. 1A). Likewise, no difference was seen in moribund fungal burden in five organs (brain, liver, spleen, kidney, lung) from randomly sampled mice from both groups (Fig. 1B). Histopathologic analysis of animals by the Duke Research Animal Pathology Service revealed vascular, multifocal dissemination of fungal elements in organs after both routes of infection (Fig. 1C).

**Figure 1.**
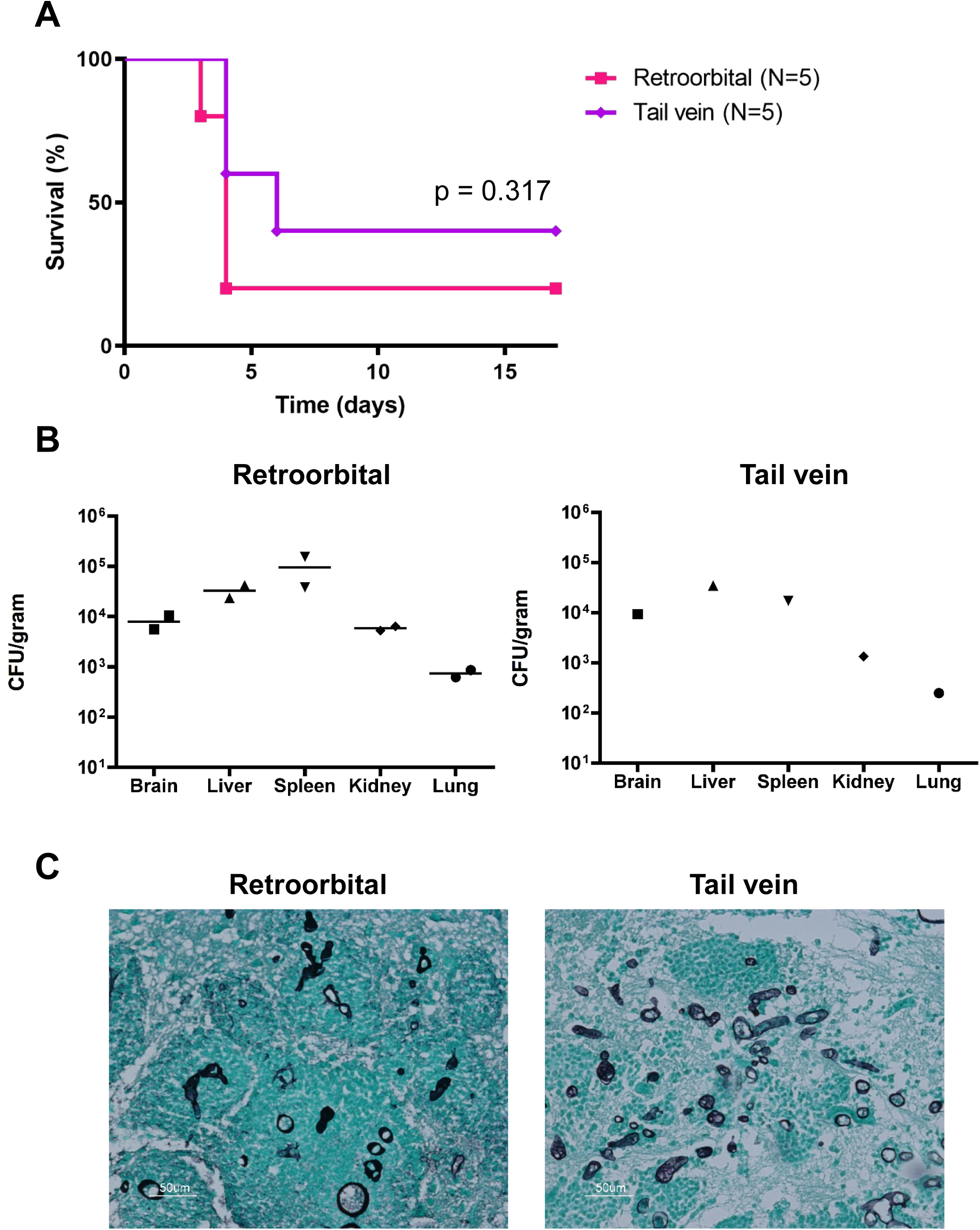
Retroorbital and tail vein injections produce equivalent outcomes. BALB/c mice were infected with 1.25 × 10^6^ spores of *M. circinelloides* f. *circinelloides* strain 1006PhL through either the retroorbital or tail vein routes. N = 5 mice per group. **A.** The two methods of infection produce equivalent mortality. **B.** The two methods of infection produce equivalent homogenized fungal burdens in randomly sampled moribund mice (N = 2 for retroorbital group, N = 1 for tail vein group). **C.** Histopathological analysis of brains from retroorbital and tail vein mice, stained with GMS (Gomori Methenamine-Silver) reveals abundant fungal elements.

### Epimutants are largely stable and revert in an organ-specific fashion during murine infection

To examine the role epimutation might play in infection, strains were recovered after one *in vivo* passage through a murine model. Immunocompetent mice were infected retroorbitally with either wild-type *Mucor* strain 1006PhL, or the *fkbA* epimutant strain SCV522. After four days, the moribund mice were euthanized with CO_2_ and five organs (brain, liver, spleen, kidney, and lung) were dissected and homogenized. Organ homogenates were plated on nonselective YPD + antibiotic agar; plates were incubated at room temperature under hypoxic conditions, which drives *Mucor* growth into discrete yeast colonies instead of hyphae, to facilitate enumeration of CFU (27). In addition, the spores of SCV522 and 1006PhL that were used to infect the mice were also plated on nonselective YPD + antibiotic agar under the same hypoxic conditions. After two days, the resulting yeast colonies were removed from hypoxia and patched onto YPD and YPD + FK506 media to test for FK506 resistance. Patched plates were grown under standard room conditions, in which normoxic exposure would lead to the resumption of hyphal growth.

The plates were imaged after 16 hours of growth and FK506-resistant (hyphal) versus FK506-sensitive (yeast) colonies were counted. All colonies grew as hyphae on nonselective YPD under normoxic conditions, as expected. Among strains isolated from the mouse infected with *fkbA* epimutant SCV522, it was noted that the epimutant strain was largely stable *in vivo*. Reversion to FK506 sensitivity was found more often in strains recovered from the brain compared to other organs (Fig. 2A). When quantified, more than half of the brain isolates had reverted to FK506 sensitivity, while reversion in the other organs was much more limited; reversion was not observed in colonies derived from the spores of SCV522 that did not undergo *in vivo* passage. In contrast, regardless of the organ, none of the isolates recovered from the 1006PhL-infected mouse were able to acquire FK506 resistance over the short time period of incubation (Fig 2B). Quantification of FK506 resistance was performed with a target number of 150 patched colonies from each organ condition; fewer colonies were used if less than 150 colonies were recovered from the organ homogenate (Table S1).

**Figure 2.**
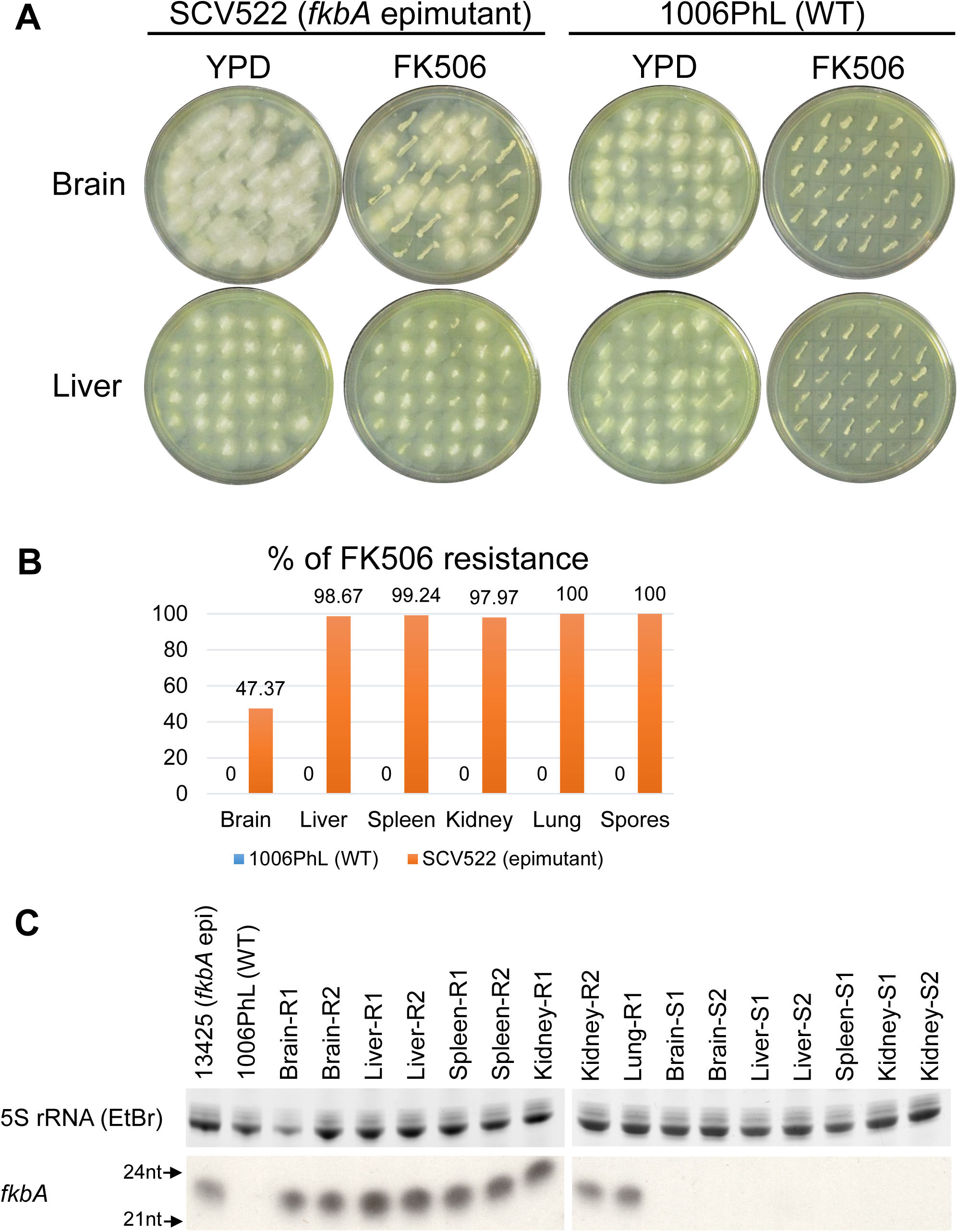
FK506 epimutants revert in an organ-specific manner after *in vivo* passage. *fkbA* epimutant and wild-type *M. circinelloides* f. *circinelloides* strains were recovered from multiple organs after *in vivo* infection/passage in one mouse each. Strains were recovered after mouse was moribund, four days post infection. SCV522, *fkbA* epimutant; 1006PhL, wild-type strain; R, resistant strain; S, sensitive strain. **A**. Representative images of colonies recovered from the brains and livers of mice infected with either SCV522 or 1006PhL and patched onto YPD with and without FK506. All colonies patched on nonselective YPD media grow as hyphae. FK506-sensitive strains patched on YPD+FK506 media grow as smaller yeast colonies. Loss of FK506 resistance from the epimutant strain is observed in the brain but not in the liver. **B**. Quantification of the percentage of FK506 resistance by organ, after *in vivo* passage. N = 150 colonies per organ; however, fewer than 150 colonies were recovered from some organs (Table S1). Spores: SCV522 and 1006PhL spores plated without *in vivo* passage. FK506 resistance was below the limit of detection in all strains derived from wild-type 1006PhL infection. **C**. sRNA hybridization of representative sensitive and resistant strains randomly selected from SCV522-infected mouse after passage. All resistant isolates continue to express sRNA against *fkbA*, whereas reverted, FK506-sensitive isolates do not. 5S rRNA was stained with ethidium bromide and served as the loading control.

Representative sensitive and resistant isolates were selected for further analysis from the strains recovered from organs after SCV522 infection. Small RNA (sRNA) hybridization revealed that all resistant strains expressed sRNAs against the *fkbA* gene, as would be expected if they had maintained their resistance through epimutation (Fig 2C). Strains which had reverted to sensitivity, regardless of organ of origin, no longer expressed sRNAs against *fkbA* (Fig 2C).

This brain-specific tendency toward reversion is also seen in an alternative *Mucor* model, in which the mice are immunosuppressed with cyclophosphamide two days prior to infection. In this model, the mice are euthanized after two days due to a more rapid course of infection; however, the brain-specific reversion of epimutation is maintained (Fig S1A, C). Similarly, analysis of representative sensitive and resistant isolates from all organs revealed that the phenotypic loss of FK506 resistance was correlated with the cessation of sRNA expression against *fkbA* (Fig S1B). Infection with 1006PhL in this model also does not lead to the development of FK506 resistance from the recovered wild-type isolates.

### *In vivo* passage causes development of *fkbA* epimutants in an organ-specific fashion after prolonged FK506 exposure

*In vivo* passage in mice without drug exposure does not lead to direct induction of epimutation-based FK506 resistance. However, passage through a murine host does lead to increased rates of epimutation if recovered strains are exposed to FK506 for longer periods of time.

As in previous experiments, immunocompetent mice were retroorbitally infected with wild-type 1006PhL spores and euthanized after four days when moribund. Organ homogenates from three mice were plated directly on YPD + antibiotics + FK506, as were 1006PhL spores that did not undergo *in vivo* passage. Plates were incubated at room temperature with light for five days; during this growth period sensitive yeast colonies arose, some of which transitioned to form resistant hyphal patches. Yeast and hyphal colonies from each organ were counted to calculate the percentage of FK506 resistance arising over time. Intriguingly, the most significant effect was once again seen with isolates from the brain (Fig 3A). In this case, brain-derived isolates developed FK506 resistance at an average rate of 6.5%, higher than the rate observed in isolates from any other organ or from spores which did not undergo *in vivo* passage.

**Figure 3.**
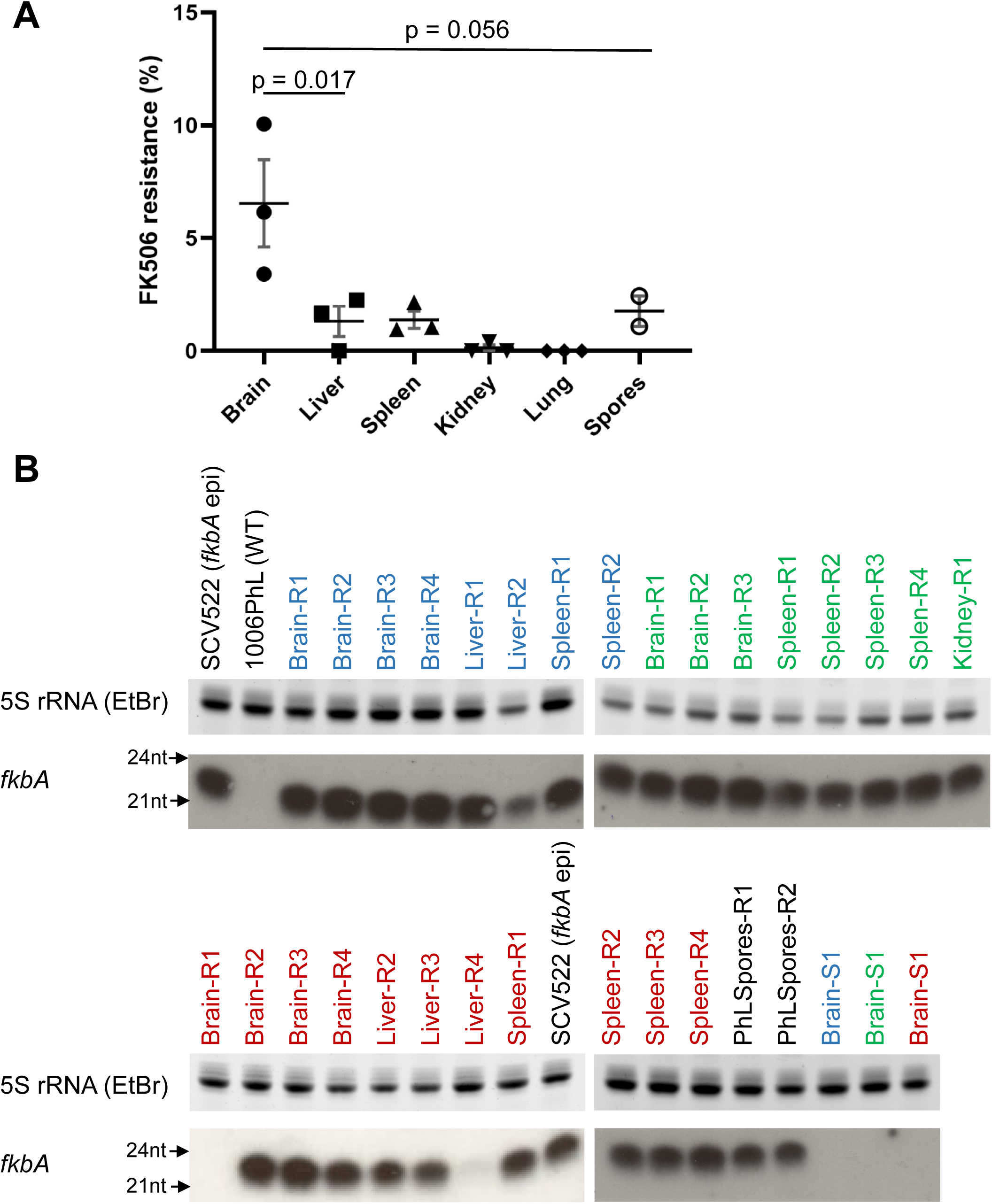
Organ-specific development of FK506 resistance after *in vivo* passage. Mice (N = 3) were infected with the wild-type 1006PhL strain; colonies were recovered on YPD+FK506 media when mice were moribund, four days post infection. Five days after recovery, resistant and sensitive colonies were counted. **A.** Percentage of FK506 resistance by organ, showing higher rates of resistance in the brain. 1006PhL, 1006Phl spores plated without *in vivo* passage. Significance determined via one-way ANOVA (P= 0.0035) with post-hoc Tukey’s Multiple Comparison test. **B**. sRNA hybridization analysis against *fkbA* demonstrates the development of sRNA expression against *fkbA* in strains from all organs and spore conditions. SCV522, *fkbA* epimutant; 1006PhL, wild-type strain; R, resistant strain; S, sensitive strain. Strains isolated from each individual mouse are identified by the color of the label (blue, green, or red). Two of the 27 resistant strains and all three of the sensitive strains (one from each mouse) show no evidence of sRNA expression against *fkbA.*

sRNA hybridization was performed to detect *fkbA* sRNA expression in representative FK506-sensitive and -resistant strains from each mouse (Fig 3B). More than 90% (25 of 27) of the resistant strains express sRNA against *fkbA,* suggesting epimutation is the predominant method by which resistance is gained after *in vivo* passage. The overall rate of FK506 resistance was much lower in strains derived from the liver, spleen, kidney, or lung, or in spores which did not undergo *in vivo* passage; however, the predominant mechanism of resistance in these isolates was still sRNA-based. Sensitive strains isolated from the brain of each mouse did not express sRNA against *fkbA* (Fig 3B).

## DISCUSSION

The phenomenon of epimutation was recently characterized as a novel mechanism of RNAi-dependent drug resistance in *Mucor* (17-19). Here, we report the first analysis of the impact of epimutation during infection. We identified two contrasting epimutant-related phenomena that occur during a murine *in vivo* model of systemic mucormycosis. First, infection with an epimutant strain demonstrated that epimutation is maintained *in vivo* over the course of infection, in multiple organs. Reversion of epimutation was markedly higher in the brains of mice than in other organs tested, at about 50%; this incomplete reversion may represent a bet-hedging strategy, enabling the fungus to respond to a range of environments. Second, *in vivo* passage of a wild-type strain of *Mucor* can lead to increased induction of epimutant-driven drug resistance, especially passage through the brain. This induction of epimutation does not appear to occur during the infection itself, but instead arises several days after recovery of strains from organs under exposure to drug selection.

It is intriguing that growth in the brain appears to induce rapid changes in epigenetic states, including both reversion and induction of epimutation-driven FK506 resistance under different circumstances. One possibility is that there is simply more fungal replication in the brain, enabling more opportunities for stochastic toggling of epigenetic states. Clinically speaking, rhino-orbito-cerebral mucormycosis is the most common manifestation of disease, and this could reflect an aspect of the cerebral environment that makes the brain especially favorable for *Mucor* growth (2, 3). Histopathologic analysis of the systemic *Mucor* infections carried out in this study correlates with clinical evidence, as qualitatively more hyphal elements were visualized in the brain than in the other organs studied (Fig 1C). Alternatively, instead of merely serving as a location for growth, the brain environment could provide a key, unidentified facet that induces epimutant switching. The rapid reversion effect was not replicated *in vitro* using commercially available media made from specific organs (tested using the bacteria media Brain Heart Infusion vs. Liver Broth; Fig S2). We note that the presence or absence of cyclophosphamide immunosuppression in the mouse model did not alter the organ-specific nature of this reversion.

Another interesting observation in this study is that induction of epimutants in the brain was not immediate. Direct plating of wild-type strains recovered from the brain did not identify any strains with FK506 resistance; conversely, resistance arose only after several days of exposure to the antifungal once strains had been recovered from organs. This suggests that the brain environment is not directly inducing *fkbA* epimutants, but perhaps priming isolates in a manner which enables them to subsequently respond rapidly to stresses including antifungal exposure. Over 90% of the resistant strains identified in this study were epimutants, which correlated with previously documented high rates of *fkbA* epimutant generation in the 1006PhL strain (17). Additional studies will need to be conducted to further examine induction of epimutation after *in vivo* exposure; it would be of interest to determine whether the rate of epimutation is increased in infected animals treated with clinical antifungals.

This is the first example in which a stressful environmental condition – exposure to the host brain – has led to increased rates of epimutation in wild-type *Mucor*. Previously tested conditions which did not increase rates of epimutation included prior epimutation (induction of a second instance of epimutation from prior epimutant strains which were reverted to wild type), antifungal drug exposure, cell wall stress, low nitrogen, low glucose, oxidative stress, and trisporic acid (mating conditions). The only described method which had increased the rate of epimutation was mutation of components of the core RNAi pathway (17). We have unsuccessfully attempted to replicate this brain-specific induction by testing various pH and hypoxia conditions (Z. Chang, unpublished data); additional study is required to elucidate the factor(s) that lead to this increased rate of resistance.

Given that *Mucor* is an environmentally ubiquitous fungus encountered on a global scale (28), it is also worthwhile to consider how non-host-pathogen interactions in the soil environment might affect traits such as epimutation. Epimutation is not limited to generating resistance to FK506 alone, but also functions to generate resistance to antifungals with entirely different mechanisms of action, such as the nucleotide analog 5-fluoroorotic acid (5-FOA) (18). Furthermore, antifungals including FK506 and rapamycin, both of which bind to FKBP12, are known to be produced by members of the bacterial genus *Streptomyces*, which are found in soil (29, 30). We speculate that exposure to these and other antifungal compounds in the natural soil environment might lead to the development of clinically relevant epimutations similar to the *fkbA* epimutation, perhaps playing a role in the intrinsic resistance of *Mucor* to many antifungal drugs. These environmental epimutations could then be rapidly induced or reverted over the course of host infection, potentially serving as virulence factors that affect treatment and clinical outcomes.

In summary, this study of epimutation revealed that epimutations affecting antifungal resistance can be both induced and reverted over the course of *in vivo* infection of *M. circinelloides*, with increased effects in the brain. Rapid switching of epigenetic states may enable adaptation to the very different environments in which *Mucor* can be found, from soil to invertebrate to vertebrate hosts. A better understanding of the mechanisms by which fungi can adapt to antifungal stress will improve our understanding of fungal biology and pathogenesis, while opening potential avenues for clinical treatment.

## MATERIALS AND METHODS

### Growth of strains

The *M. circinelloides* f. *circinelloides* strain 1006PhL served as the wild type for all studies. *fkbA* epimutant strain SCV522 was derived from 1006PhL in a prior study, and recovered from −80°C laboratory freezer stocks (17). All strains were grown at room temperature (approximately 24°C) with normal daylight exposure. Strains were cultured on YPD agar (10 g/L yeast extract, 20 g/L peptone, 20 g/L dextrose). The antifungal drug FK506 (Prograf, in a sterile formulation from Astellas Pharma) was sterilely added to YPD media as required after autoclaving, to achieve a final concentration of 1 μg/mL.

### Murine infection model

Mice used in this study were male BALB/c mice (Charles River) at 6 weeks of age, weighing approximately 20 to 25 grams. Mice were infected intravenously with a dose of 1.25 × 10^6^ spores of 1006PhL suspended in 100 μL of sterile phosphate buffered saline (PBS). Infected mice were monitored by weighing and visual scoring twice daily, and euthanized via CO_2_ exposure when humane endpoints were reached (including but not limited to weight loss greater than 20% of starting body weight, or significant loss of grooming or mobility).

All intravenous infections were performed via retroorbital injection unless otherwise noted, an approach developed based on previously published protocols (23). To perform retroorbital injections, mice were anesthetized by exposure to inhaled isoflurane. Fungal inoculum suspended in PBS was then injected behind either the right or left eye, and the animal was visually monitored while recovering from anesthesia in room air.

As needed, mice were immunosuppressed two days prior to fungal infection. A single dose of 200 mg/kg cyclophosphamide in sterile PBS was delivered by intraperitoneal injection.

### Strain isolation after *in vivo* passage

Mice were euthanized via CO_2_ exposure at the humane endpoint, and up to five organs (brain, liver, spleen, kidney, lung) were dissected. Whole organs were homogenized in 1 mL of sterile PBS through bead beating in a tube with two 5 mm steel beads (3 cycles of 1 minute each). Organ homogenates were plated on YPD agar containing antibiotics (50 μg/ml ampicillin and 30 μg/ml chloramphenicol) for selective isolation of fungal colonies.

When hypoxic conditions were required, the GasPak EZ Container System was used (BD Diagnostics). YPD + antibiotic plates were placed in the GasPak Large Incubation Chamber and three anaerobe satchets were added prior to sealing the chamber. Maintenance of hypoxic conditions (less than 1% O_2_ and greater than 13% CO_2_) was monitored using Dry Anaerobic Indicator Strips (BD).

If FK506 selection was required, 1 μg/mL FK506 was added to the YPD + antibiotic agar. Representative FK506-resistant (hyphal) and sensitive (yeast) colonies were picked for further analysis.

### Fungal burden

Fungal burden was measured by plating organ homogenates (as described above) on YPD + antibiotic agar and counting colonies. This method does not produce a quantitative total for fungal burden in organs, due to the breakup of hyphae into fragments during the bead beating step; however, the resulting counts allow for qualitative comparison between test groups of a given experiment and recovery of isolates for further analysis.

### sRNA extraction and hybridization

Strains for sRNA extraction were grown on plates overlaid with sterile cellulose film (ultraviolet irradiated for 10 minutes per side) to allow for easier removal of hyphae. Small RNAs were extracted from hyphae using the mirVana miRNA Isolation kit (Ambion, Foster City, CA). 3.5 μg of purified sRNA for each sample was separated by electrophoresis on 15% TRIS-urea gels.

When the 5S rRNA loading control was analyzed by ethidium bromide staining, sRNA gels were cut in half prior to transfer; the top half was stained with ethidium bromide and visualized under UV light, while the bottom half was transferred to Hybond N+ filters, and cross-linked by ultraviolet irradiation (2 pulses at 1.2 × 10^5^ μJ per cm^2^) (17). When the 5S rRNA loading control was analyzed by radioactive probing instead, the entire gel was transferred to the membrane as described.

Prehybridization of membranes was carried out using UltraHyb buffer (Ambion) at 65°C. *fkbA* antisense-specific and 5S rRNA riboprobes were prepared by *in vitro* transcription using the Maxiscript kit (Ambion); primers are listed in Table S2. Riboprobes were treated by alkaline hydrolysis as previously described (31), to generate an average final probe size of ∼50 nucleotides. Hybridization was carried out overnight.

### Statistics

Statistical significance of the Kaplan-Meier survival curves was determined using a log-rank test. One-way ANOVAs were used to determine significance when comparing rates of FK506 resistance, with Tukey’s Multiple Comparison Test as a post-hoc test where appropriate. All statistical analysis was performed using GraphPad Prism.

## ACKNOWLEDGEMENTS

We thank Shelby Priest for critical reading of the manuscript. We would also like to thank Dr. Jeffery Everitt, DVM of the Duke Research Animal Pathology Service for histopathologic analysis.

## SUPPLEMENTAL MATERIAL

**Figure S1.**
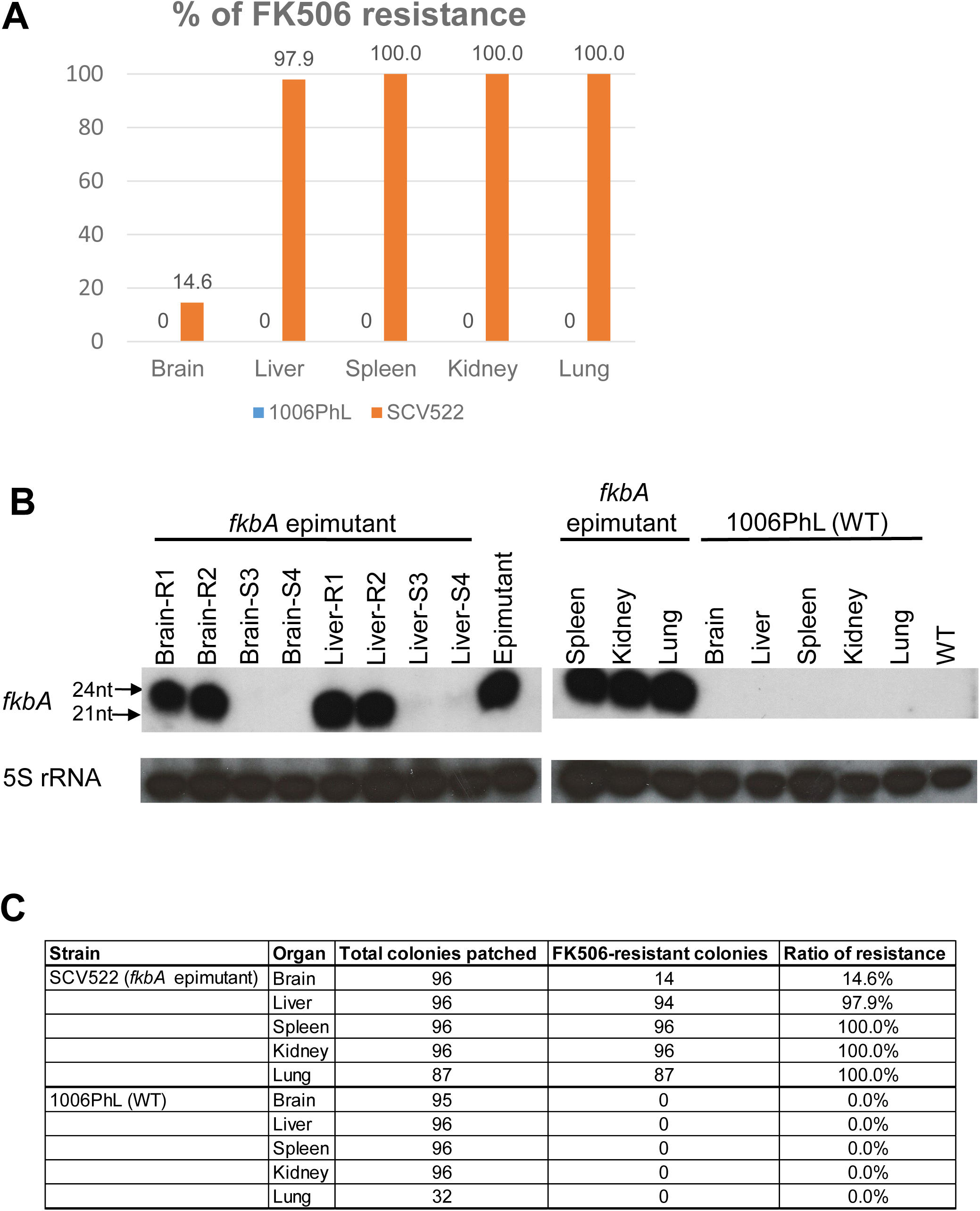
Organ-specific epimutant reversion is also observed in an immunosuppressed murine model. Brain-specific reversion was also observed after passage of strain SCV522 through a BALB/c mouse immunosuppressed with cyclophosphamide. Strains were recovered when the animal was moribund, two days post infection. **A.** Quantification of the percentages of FK506 resistance by organ. FK506 resistance was below the limit of detection in all strains derived from wild-type 1006PhL infection. **B.** Representative sensitive and resistant colonies from the brain and liver were selected for sRNA hybridization analysis, along with resistant colonies from the spleen, kidney, and lung. sRNA expression against *fkbA* is observed in resistant but not sensitive colonies. *fkbA* and 5S rRNA loading control were quantified using radioactive probes. **C.** Table of colony counts from which ratios in panel B were derived.

**Figure S2.**
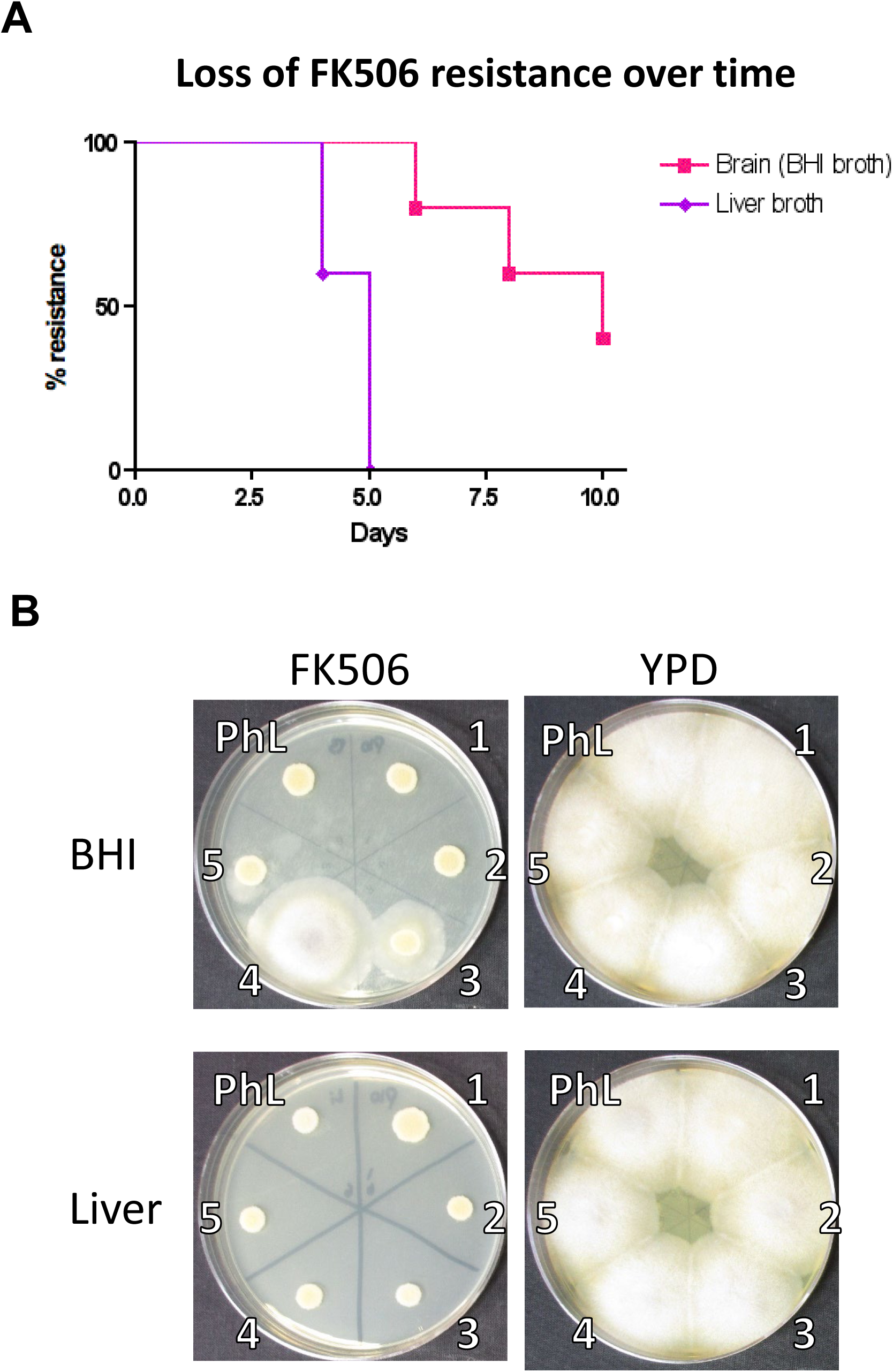
Organ-specific epimutant reversion is not replicated in vitro with organ-based media. Strains of epimutant SCV522 or wild-type 1006PhL were passaged daily for 10 passages in commercially available organ-based liquid medias. BHI, Brain-Heart Infusion; Liver, Liver broth. **A.** Curve showing the loss of FK506 resistance in strains passed on each media type, over time. N = 5 independent passages per media. In contrast to the *in vivo* brain reversion effect, reversion was seen more rapidly after passage in liver broth compared to BHI broth. **B.** Phenotypic analysis of all passaged strains after the 10^th^ passage, grown on YPD and YPD + FK506 media. BHI, strains which were passed on BHI media. Liver, strains which were passaged on liver media. PhL, 1006PhL strain after 10 passages; strains 1-5 are independent passages of epimutant strain SCV522 which were passed on either BHI or Liver media, as labeled. All epimutant strains passed on liver broth have reverted to FK506 sensitivity after 10 passages, while some epimutant strains passed on BHI have maintained their resistance.

**Table S1.** Numbers of sensitive and resistant colonies for Figure 2B. Table showing colony numbers from which ratios in Figure 2B were derived. Target N for all conditions was 150 colonies; however, fewer than 150 colonies were recovered from some organs.

| Strain | Organ | Total colonies patched | FK506-resistant colonies | Ratio of resistance |
| --- | --- | --- | --- | --- |
| SCV522 (*fkbA* epimutant) | Brain | 57 | 27 | 47.4% |
|  | Liver | 150 | 148 | 98.7% |
|  | Spleen | 132 | 131 | 99.2% |
|  | Kidney | 148 | 145 | 98.0% |
|  | Lung | 1 | 1 | 100.0% |
| 1006PhL (WT) | Brain | 88 | 0 | 0.0% |
|  | Liver | 150 | 0 | 0.0% |
|  | Spleen | 150 | 0 | 0.0% |
|  | Kidney | 156 | 0 | 0.0% |
|  | Lung | 4 | 0 | 0.0% |
| 1006PhL spores (without *in vivo* passage) | N/A | 150 | 0 | 0.0% |
| SCV522 spores (without *in vivo* passage) | N/A | 150 | 150 | 100.0% |

**Table S2.**
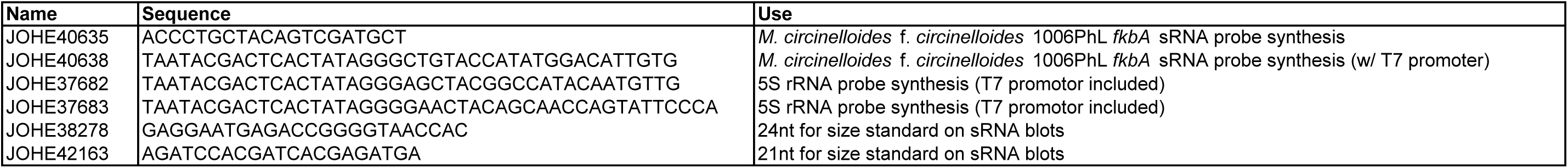
Primers used in this study.

